# Making ends meet: new functions of mRNA secondary structure

**DOI:** 10.1101/2020.04.29.069203

**Authors:** Dmitri N. Ermolenko, David Mathews

## Abstract

The 5’ cap and 3’ poly(A) tail of mRNA are known to synergistically regulate mRNA translation and stability. Recent computational and experimental studies revealed that both protein-coding and non-coding RNAs will fold with extensive intramolecular secondary structure, which will result in close distances between the sequence ends. This proximity of the ends is a sequence-independent, universal property of most RNAs. Only low-complexity sequences without guanosines are without secondary structure and exhibit end-to-end distances expected for RNA random coils. The innate proximity of RNA ends might have important biological implications that remain unexplored. In particular, the inherent compactness of mRNA might regulate translation initiation by facilitating the formation of protein complexes that bridge mRNA 5’ and 3’ ends. Additionally, the proximity of mRNA ends might mediate coupling of 3′ deadenylation to 5′ end mRNA decay.

## 1. Introduction

The sequence and secondary structures at the 5’ and 3’ termini of RNA play important roles in cellular processes (Chatterjee & Pal, 2009; Genuth & Barna, 2018; Hinnebusch, Ivanov, & Sonenberg, 2016). Both the 5’ and 3’ ends of RNA are recognized by proteins that mediate RNA processing or mRNA translation (Curry, Kotik-Kogan, Conte, & Brick, 2009). In eukaryotes, transcripts produced by RNA polymerase II are modified with by a 7-methyl-guanosine cap (m^7^Gppp cap) at the 5’ end and a poly(A) tail at the 3’ end.

Both the 5’ cap and the 3’ poly(A) protect mRNA from degradation and stimulate translation. Furthermore, the regulation of both mRNA degradation and translation involve interactions of the 5’ cap and 3’ poly(A) tail that are protein mediated. The pervasiveness across evolution of protein bridges spanning the ends of mRNA in translation regulation and decay is surprising because mRNA circularization should incur a substantial entropic cost as compared to a random coil (Yoffe, Prinsen, Gelbart, & Ben-Shaul, 2011). However, new studies discussed below suggest that the expected entropic penalty is mitigated by intramolecular basepairing interactions that provide the energetic drive for compaction. The realization that mRNA ends are intrinsically close may have many important mechanistic and evolutionary implications that await further investigation.

## 2. RNA HAS THE INTRINSIC PROPENSITY TO FOLD INTO STRUCTURES THAT BRING THE SEQUENCE ENDS CLOSE

### 2.1. The formation of intramolecular RNA secondary structure brings the ends into proximity

It has long been known that nucleotides adjacent to the 5’ end are basepaired with nucleotides adjacent to the 3’ end in a number of non-coding RNA molecules, such as tRNA, 5S rRNA, 23S rRNA and the RNA component of RNase P (Fox & Woese, 1975; Gutell, Gray, & Schnare, 1993; Holley et al., 1965; James, Olsen, Liu, & Pace, 1988). The basepairing between the ends in these RNAs is rather amazing considering significant variations in RNA length, from 76 (tRNA) to 2900 nucleotides (23S rRNA). As more RNA structures were determined over the years, the list of different RNA molecules showing basepairing between the 5’ and 3’ ends continued to grow (Vicens, Kieft, & Rissland, 2018). However, this common feature of many RNA structures attracted little attention from investigators.

Recent computational studies, focusing on RNA secondary structure formation, suggested that the ends of RNA sequences are close in space, regardless of sequence composition and length (Clote, Ponty, & Steyaert, 2012; Fang, 2011; Yoffe et al., 2011). Yoffe et al. estimated that the average 5’ to 3’ end distance in RNAs is 3 nm (Yoffe et al., 2011). The proximity of RNA ends arises naturally from stem-loop formation. Helices shorten the end-to-end distance and, with increasing helix formation, the probability increases that nucleotides at the 5’ end will be basepaired to nucleotides at the 3’ end (Fig. 1a-b). It is known from prior studies that even random sequences (composed of all four base identities) will form extensive secondary structure (J. H. Chen, Le, Shapiro, Currey, & Maizel, 1990; Clote, Ferre, Kranakis, & Krizanc, 2005; S. V. Le, Chen, Currey, & Maizel, 1988; S. Y. Le, Chen, & Maizel, 1989; Uzilov, Keegan, & Mathews, 2006; Workman & Krogh, 1999).

**Figure 1.**
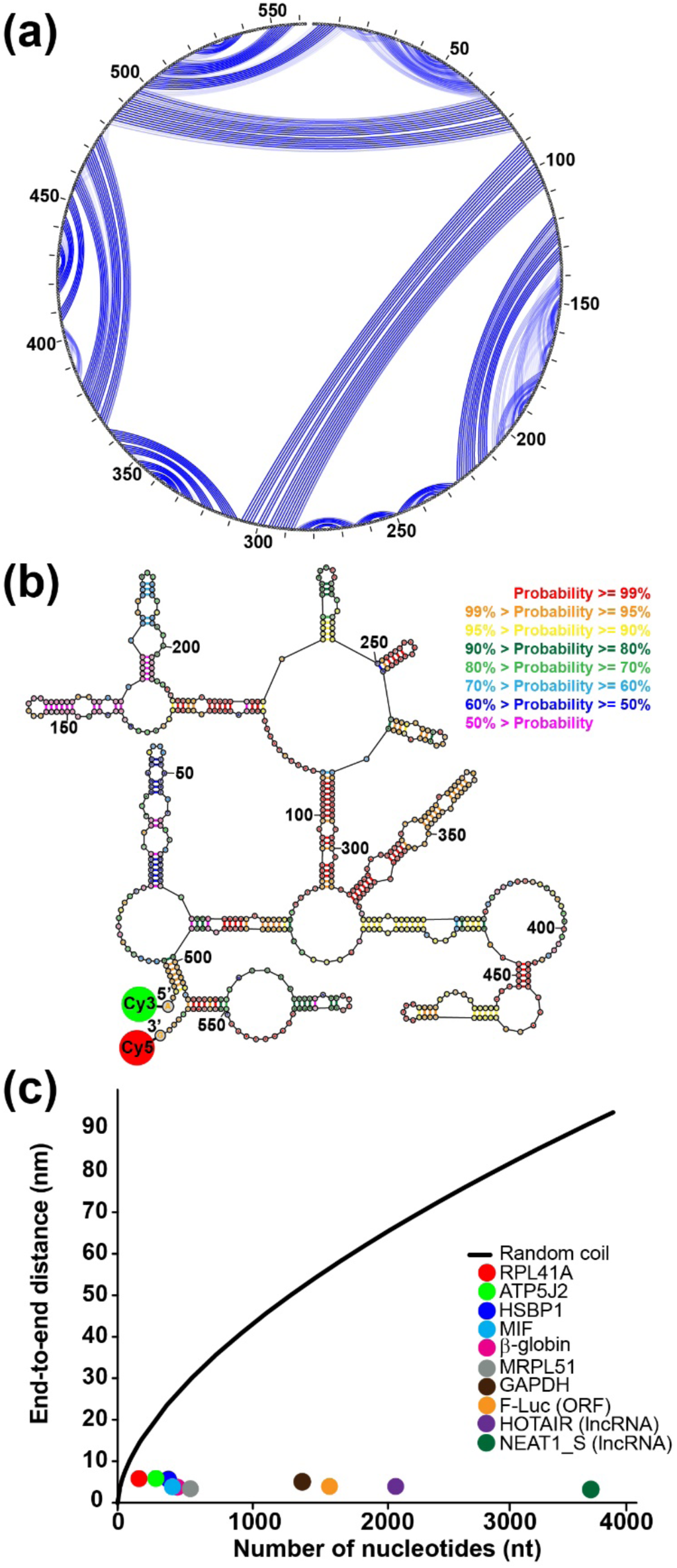
Intramolecular basepairing brings the ends of RNA close. Panels **a** and **b** show the secondary structure of the human MIF mRNA as predicted and drawn using the RNAstructure software package (*https://rna.urmc.rochester.edu/RNAstructure.html*). Panel **a** shows a circular diagram where the sequence is clockwise around the outside of the circle, with the 5’ and 3’ ends at the top of the circle. Blue lines are basepairs; the weight of a blue line represents the estimated pairing probability in the Boltzmann ensemble, where heavier lines are higher estimated probabilities. Panel **b** shows a collapsed diagram of one secondary structure in the ensemble, where basepairs are colored according to estimated base pairing probabilities in the conformational ensemble. Both representations of the secondary structure show how basepairing brings the ends close. The probable helix close to the 5’ end and the probable stem-loop at the 3’ end both serve to bring the ends together for this sequence. Panel **c** shows the FRET-measured end-to-end distances as a function of sequence length. The colored dots are: yeast RPL41A mRNA (red), firefly luciferase mRNA (orange), rabbit β-globin mRNA (magenta), human ATP5J2 mRNA (green), HSBP1 mRNA (indigo), MIF mRNA (blue), MRPL51 mRNA (grey), GAPDH mRNA (brown), HOTAIR lncRNA (purple), and NEAT1_S lncRNA (dark green). The black line is the end-to-end distance of a freely jointed RNA chain. This figure is reproduced from (Lai et al., 2018).

These computational predictions were first tested using several viral RNAs and mRNAs from the fungus *Trichoderma atroviride* with lengths from 500 to 5,000 nucleotides (Leija-Martinez et al., 2014). These sequences were folded *in vitro* without any protein factors, and single-molecule Förster resonance energy transfer (smFRET) was observed between the two ends of the molecules. When FRET was detected, the end-to-end distance ranged from 5 to 9 nanometers (Leija-Martinez et al., 2014). These results are consistent with the computational studies suggesting that the 5’ and 3’ ends in all RNAs are invariably close and tend to basepair to each other.

More recently, we further tested this hypothesis by measuring FRET between donor and acceptor fluorophores introduced at the 5’ end of 5’ UTR and 3’ end of 3’ UTR, respectively, in eight yeast and human mRNAs (Fig. 1b) (Lai et al., 2018). We measured ensemble FRET in doubly-labelled mRNA molecules, folded *in vitro* in the absence of proteins, and FRET was observed for all eight tested mRNAs. The average end-to-end distances, determined for each transcript from ensemble FRET data, were between 5 and 7 nm. These distances are independent of sequence length (Fig. 1c) and these distances are up to ten times shorter than those predicted by the freely jointed chain model for RNA random coils (Cantor & Schimmel, 1980; Grosberg & Khokhlov, 1994) (Fig. 1c). In addition, FRET between fluorophores attached to the 5’ and 3’ ends was detected in two well-studied long non-coding (lnc)RNAs, HOTAIR and NEAT1_S (Fig. 1c), providing additional support for the hypothesis about universal closeness of RNA ends (Lai et al., 2018).

Introduction of unstructured sequences, such as CA repeats, into the 5’ and 3’ UTRs of an mRNA led to disappearance of FRET between fluorophores attached to the 5’ end of the 5’ UTR and 3’ end of the 3’ UTR (Lai et al., 2018). These results indicated that the 5’ and 3’ ends of the wild-type mRNA sequence were brought within FRET distance via the formation of intramolecular basepairing. Additional FRET experiments also showed that the poly(A) tail is not involved in basepairing interactions with the 5’ UTR (Lai et al., 2018).

smFRET measurements in individual mRNA molecules revealed that instead of folding into one stable structure, mRNAs fold into an ensemble of several structural states with distinct end-to-end distances (Fig. 2) (Lai et al., 2018) and the strands interconvert between these structures. The spontaneous interconversion as observed by smFRET traces and demonstrated rates similar to those previously measured for spontaneous transitions between two alternative 5 basepair-long RNA helixes (Furtig et al., 2007) (Fig. 2). These data indicated that analogous structural rearrangements spontaneously occur in mRNAs.

**Figure 2.**
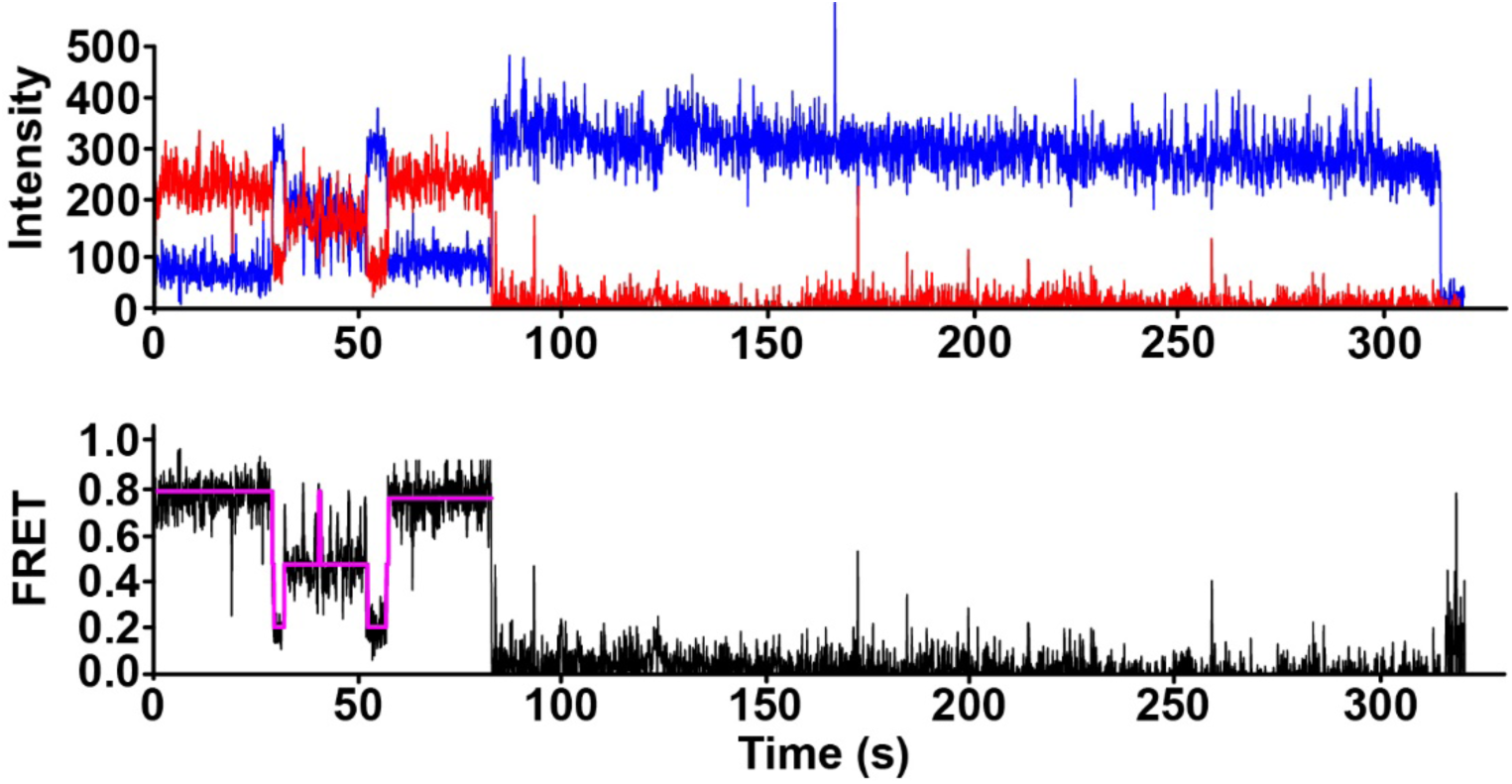
smFRET shows that mRNAs fold into a dynamic ensemble of structures. The top plot shows the smFRET for the human GAPDH mRNA as a function of time where blue is fluorescence intensity of the donor (Cy3) at the 3’ end of the 3’ UTR and red is fluorescence intensity of the acceptor (Cy5) at the 5’ end of the 5’ UTR. The acceptor photobleaches at about 80 seconds. The bottom plot shows the FRET efficiency in black with an idealized model fit by a Hidden Markon Model in magenta, where fluctuation is shown between the 0.2, 0.4, and 0.8 FRET states that correspond to distinct end-to-end distances. This figure is reproduced from (Lai et al., 2018).

### 2.2. Most mRNA and lncRNA sequences have the propensity to fold into structures with short end-to-end distances

We developed new software within *RNAstructure* for modeling the distribution of end-to-end distances of conformational ensembles (Lai et al., 2018). The ETEcalculator program estimates the end-to-end (ETE) distance for an input RNA sequence. ETE calculator samples structures from the Boltzmann ensemble (Ding & Lawrence, 2003) and estimates the end-to-end distance for each structure using polymer theory (Aalberts & Nandagopal, 2010). The estimated end-to-end distance is the mean distance across the sampled structures as reported in nm. The mean end-to-end distances from this software correlate with our ensemble FRET measurements.

To evolve sequences to have a longer end-to-end distance, we wrote the orega (“optimize RNA ends with a genetic algorithm”) program. This is also part of the RNAstructure software package, available at http://rna.urmc.rochester.edu. Orega uses a genetic algorithm to randomly evolve a region of an input sequence to maximize its fitness. Fitness is defined as the mean probability that nucleotides are unpaired in the region, as estimated by secondary structure prediction (Mathews, 2004), plus the linguistic complexity in the region. The linguistic complexity quantifies the sequence diversity, where larger values (bounded by 1) indicate that the sequence has little repetition, and small values (bounded by 0) indicate sequence repeats (Gabrielian & Bolshoy, 1999; Trifonov, 1990). We found the complexity was an important aspect to evolve sequences that could be successfully cloned.

We applied the software to make estimates for the distances between the 5’ end of 5’ UTR and 3’ end of 3’UTR across the HeLa human cell transcriptome of ∼21,000 transcripts. The estimated distances were relatively narrowly distributed with a mean of ∼4 nm (Fig. 3a). It was rare to have a long end-to-end distance (defined as longer than 8 nm); about ∼0.01% of mRNAs were estimated to have a long end-to-end distance (Lai et al., 2018). Likewise, the estimated end-to-end distances in ∼104,000 human RNA sequences annotated as lncRNAs were relatively narrowly distributed with a mean of ∼4 nm (Fig. 3b). Only ∼0.12 % of all lncRNAs were predicted to have a long end-to-end distance (Lai et al., 2018). Hence, the intrinsic propensity of RNA structure to result in short end-to-end distances appear to be common to all human mRNAs and lncRNAs. Furthermore, the proximity of RNA ends appears to be largely independent of sequence and length.

**Figure 3.**
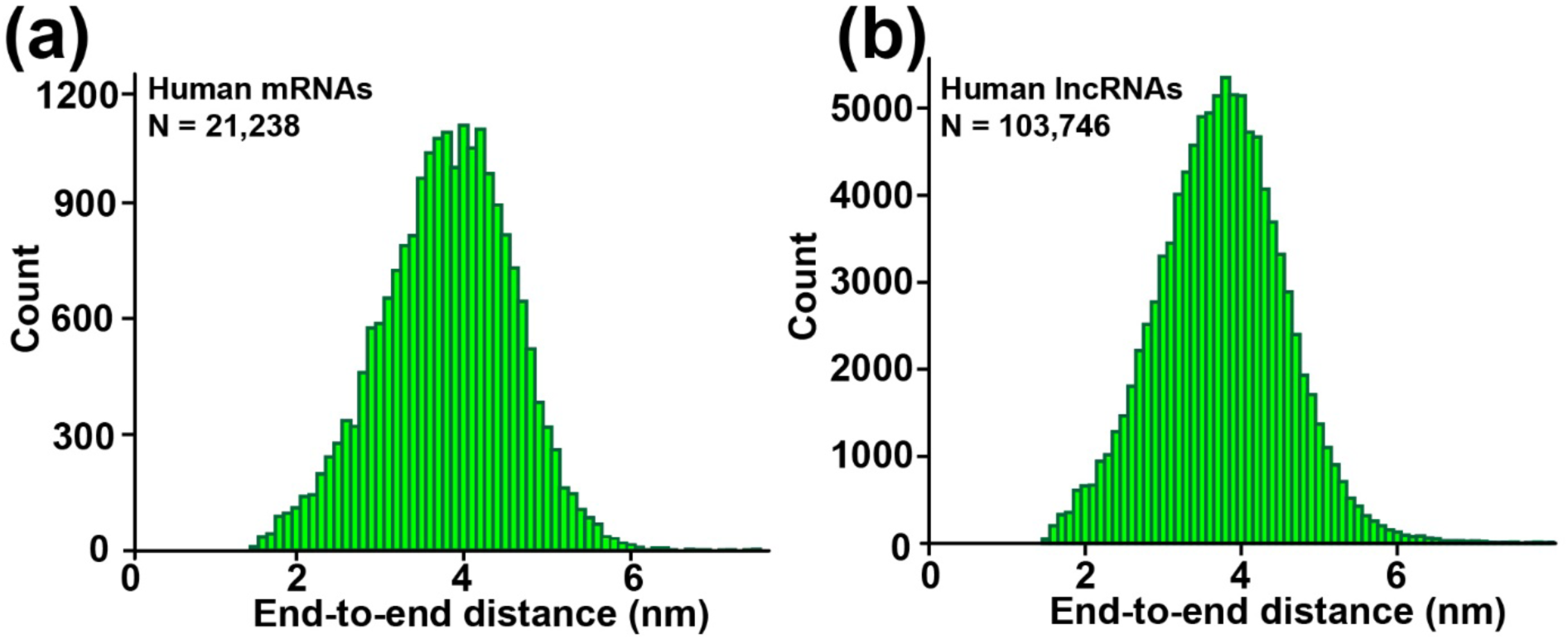
Histogram of end-to-end distances for human mRNAs and lncRNAs. Panel **a** is the distribution of estimated mRNA end-to-end distances for the HeLa cell transcriptome. Panel **b** is the distribution for human lncRNA sequences. N is the number of sequences analyzed. This figure is reproduced from (Lai et al., 2018).

### 2.3. mRNA 5’ and 3’ UTRs basepair *in vivo*

The intrinsically close end-to-end distances in mRNA and lncRNA are likely also found in live cells. mRNA secondary structure might be disrupted by helicases and other RNA binding proteins. Furthermore, the ribosome efficiently unwinds the secondary structure of mRNA within an Open Reading Frame (ORF) by translocating along the mRNA during the elongation of the polypeptide chain (Takyar, Hickerson, & Noller, 2005; Wen et al., 2008). Nevertheless, a number of structured mRNA elements are known to regulate translation initiation, including bacterial riboswitches (Roth & Breaker, 2009), frameshift-inducing hairpins and pseudoknots of eukaryotic viruses (Giedroc & Cornish, 2009), Internal Ribosome Entry Sites (IRES) (Mauger, Siegfried, & Weeks, 2013), Iron Response Elements (IRE) in the 5’ UTR of transcripts coding for proteins involved iron metabolism (Leipuviene & Theil, 2007), and Cap-Independent Translational Enhancers (CITEs) (Simon & Miller, 2013). Studies have also shown that protein binding sites on mRNA are determined by accessibility, as governed by the RNA structure (Li, Kazan, Lipshitz, & Morris, 2014; Li, Quon, Lipshitz, & Morris, 2010). Similarly, accessibility to oligonucleotide binding in siRNAs, miRNAs, and antisense oligonucleotides is also governed by the RNA structure, which can occlude sites (Z. J. Lu & Mathews, 2008a, 2008b; Shao et al., 2007; Tafer et al., 2008). Hence, mRNA secondary structure of is important in the cell in spite of the activities of helicases and other RNA-binding proteins (Aw et al., 2016; Z. Lu et al., 2016; Sharma, Sterne-Weiler, O’Hanlon, & Blencowe, 2016; Wu & Bartel, 2017).

Transcriptome-wide chemical probing studies in yeast, plant and human cells also support the importance of secondary structure *in vivo* (Aw et al., 2016; Ding et al., 2014; Z. Lu et al., 2016; Rouskin, Zubradt, Washietl, Kellis, & Weissman, 2014; Sharma et al., 2016). Intramolecular basepairing between distant segments of mRNA, including interactions between the 5’ and 3’ UTRs, were also observed by mapping of RNA-RNA interactions by psoralen cross-linking in yeast and human cells detected (Aw et al., 2016; Z. Lu et al., 2016; Sharma et al., 2016). These experimental results support the idea that the intrinsic propensity of RNAs to fold into structures with short end-to-end distances applies to RNA folding within cells.

### 2.4. Is short end-to-end distance a common feature of RNA and proteins?

“Closeness of the ends” has somewhat different meaning in the case of proteins and RNAs. We and others compare the end-to-end distance in folded RNA with the end-to-end distance expected for the random coil conformation of RNA. In this context, the closeness of mRNA (or lncRNA) ends means that the distance between the 5’ and 3’ ends is much shorter than the distance expected for the random coil conformation of the sequence. With exception of few RNAs, such as 16S rRNA, 23S rRNA, some RNA aptamers and ribozymes, most of RNA molecules including mRNAs do not fold into highly-condensed structures (Seetin & Mathews, 2011; Yoffe et al., 2008). Because of that, closeness of RNA ends is counterintuitive and is rather remarkable.

In contrast to RNA, most proteins fold into compact, globular structures. For that reason, the distance between N and C termini is usually examined in regard to dimensions of folded proteins (e.g. radius of gyration). In this context, the closeness of protein termini implies that the distance between N and C termini is significantly shorter than the distance expected based on chance and dimensions of a given protein. There is no agreement between different analyses on whether protein termini are generally closer than expected by chance or not (Christopher & Baldwin, 1996; Thornton & Sibanda, 1983). Nevertheless, similar to RNA, in most proteins, end-to-end distance is shorter in the folded state than in unfolded ensembles of states of a protein (Schuler & Eaton, 2008).

## 3. BIOLOGICAL IMPLICATIONS OF THE INTRISIC COMPACTNESS OF RNA

### 3.1. The intrinsic closeness of RNA ends as an evolutionary hurdle

High basepairing potential and intrinsic compactness of most natural RNA sequences have multiple evolutionary implications. mRNAs and RNA-binding proteins may co-evolve to overcome or, in some cases, to exploit the intrinsic closeness of mRNA ends. Because most, if not all, mRNA and lncRNA sequences have a propensity to form extensive secondary structures, RNA helicases and single-strand RNA binding proteins are required to keep RNA in the single-stranded conformation in live cells. RNA unwinding likely constitutes a significant energy expenditure for the cell. When RNA helicases and single-strand RNA binding proteins disassociate from RNA, RNA likely rapidly folds into compact structures.

The tendency of RNA ends to basepair to each other creates an evolutionary hurdle when RNA function requires one of RNA ends to form intermolecular baseparing interactions with another RNA molecule or to bind protein factors. One evolutionary strategy to overcome this problem may be favoring intrinsically unstructured sequences at one RNA end. For example, poly(A) sequences in the 5′ UTR of poxvirus mRNAs (Shirokikh & Spirin, 2008) and CAA nucleotide triplet repeats in the Ω leader (5’ UTR) of tobacco mosaic virus mRNAs (Agalarov, Sakharov, Fattakhova, Sogorin, & Spirin, 2014) facilitate the recruitment of the small ribosomal subunit and make translation initiation on these mRNAs independent of the presence of the 5’ cap on mRNA.

What are the defining properties of intrinsically unstructured sequences? To better understand the connection between sequence and end-to-end distance of RNA, as mediated by basepairing, we developed software to manipulate sequences to adjust end-to-end distances. We used a genetic algorithm, a type of *in silico* evolution algorithm (Lai et al., 2018), to randomly evolve a population of sequences that avoid intramolecular basepairing. At each step, either mutation or crossover (a combination of two sequences from the population) occurs, and then sequences with the best fitness are retained for subsequent refinement. Our fitness metric can be tailored to the goal, and the first metric was low mean basepairing probabilities for a stretch of the sequence. *In silico* evolution of the human GAPDH mRNA sequence confirmed that a reduction in average basepairing probability leads to an increased end-to-end distance of RNA (Lai et al., 2018). We also observed that in order to reduce basepairing probabilities, an RNA sequence increases the cytosine content and also nearly eliminates guanines. Guanosines form Watson-Crick G-C and also wobble G-U pairs, which have folding stabilities that are similar to Watson–Crick A-U base pairs and are nearly isosteric to A-U pairs (J. L. Chen et al., 2012; Varani & McClain, 2000). Therefore, it appears likely that sequences must avoid guanines to avoid basepairing. As sequences are evolved *in silico* to reduce the average basepairing probability, we also observe that the linguistic complexity decreases (Gabrielian & Bolshoy, 1999; Troyanskaya, Arbell, Koren, Landau, & Bolshoy, 2002). The complexity is a measure of the sequence repetition, where low complexity indicates that the sequences are repetitive (Lai et al., 2018).

Our *in silico* evolution experiments showed that guanosine depletion and low sequence complexity are necessary but not sufficient for an RNA to be unstructured (Lai et al., 2018). Therefore, specific, low-complexity sequences of adenosines, cytosines, and uracils will be random coils. Therefore, unstructured RNA sequences in organisms are likely to have resulted from intense natural selection, and theses sequences are likely serving biological roles. Transcriptome-wide search for the intrinsically unstructured RNA segments might reveal novel regulatory sequences in mRNA.

Another evolutionary strategy for keeping RNA ends apart may be the sequestration of one of the RNA ends by pseudoknotted basepairing interactions with nucleotides in the middle of the RNA. 16S rRNA is an example of this strategy. In contrast to 5S or 23S rRNAs, whose ends are basepaired, the 5’ and 3’ ends of 16S rRNA in the small ribosomal subunit are over 8 nm away from each other. Translation initiation in bacteria is facilitated by baseparing interactions between the 3’ end of 16S rRNA and the Shine-Dalgarno (ribosome-binding) sequence in mRNA (Figure 4). Residues adjacent to the 5’ end of 1542 nucleotide-long 16S rRNA (*E.coli* numbering) form intramolecular helixes h2 (a pseudoknot) and h3 by basepairing with residues 916-918 and residues 547-556, respectively (Noller & Woese, 1981; Yusupov et al., 2001). These interactions might have evolved to keep the 3’ end of 16S rRNA free to bind the Shine-Dalgarno sequence in mRNAs.

**Figure 4.**
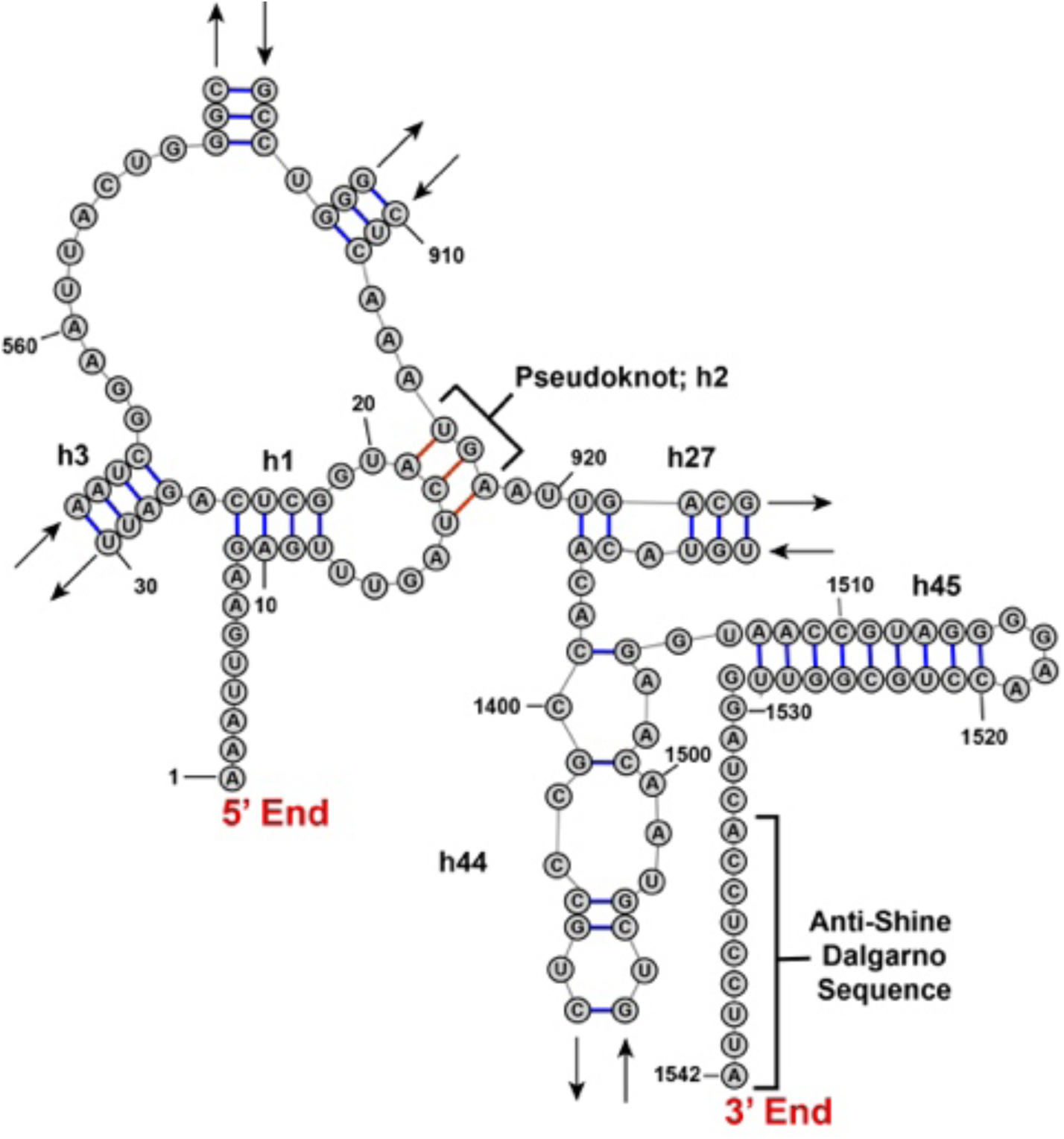
The *E. coli* 16S rRNA sequence ends are far apart. This figure shows how a pseudoknot in the small subunit rRNA facilitates a longer end-to-end distance than we found in other ncRNA. In part, this exposes the antiShine-Dalgarno sequence to base pairs with an mRNA Shine-Dalgarno sequence to initiate translation.

**Figure 5.**
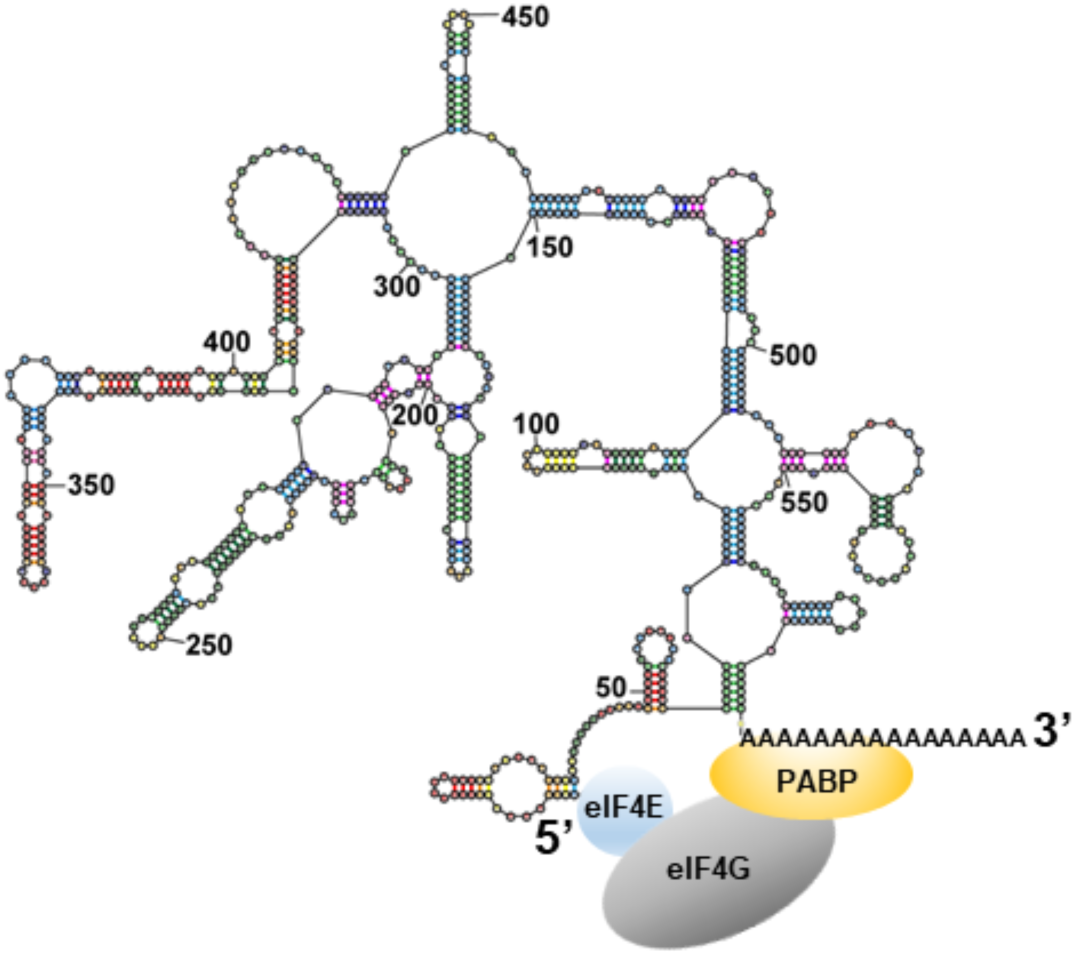
The intrinsic closeness of mRNA ends may augment translation by stabilizing the eIF4E•eIF4G•PABP complex.

### 3.2. mRNA compactness in mRNA decay

Closeness of RNA ends may not always be an evolutionary obstacle. Indeed, as discussed below, a number of RNA binding protein complexes emerged through evolution to regulate translation initiation and mRNA decay by bridging mRNA ends. Binding of these complexes to mRNA is likely stabilized by the intrinsic closeness of mRNA ends. The prominence of protein-mediated interactions between mRNA ends in evolution of protein synthesis and mRNA decay suggest that the protein complexes might have evolved to exploit the intrinsic closeness of RNA ends. Therefore, the innate compactness of mRNA likely facilitates the regulation of translation and mRNA degradation.

Protein-mediated interactions between mRNA ends are involved in regulation of mRNA degradation. Eukaryotes have two major pathways of mRNA decay: exosome-catalysed degradation from the 3’ end and 5’-to-3’ degradation. 5’-to-3’ degradation mRNA degradation begins with poly(A) tail shortening or deadenylation carried out by two multiprotein complexes, PAN2-PAN3 and CCR4-NOT. Dedadenylation is followed by the recruitment of Dcp1-Dcp2-Edc4 de-capping complex to 5’ end of mRNA, the 5’ cap removal and 5’-to-3’ exonucleolytic degradation of mRNA by Xrn1 exonuclease (Mugridge, Coller, & Gross, 2018). mRNA deadenylation and decapping are coupled through protein-protein interactions of CCR4-NOT and decapping complex (Mugridge et al., 2018). In budding and fission yeast, deadenylation is also linked to decapping by the interaction between Pat1-Lsm1-7 complex, which recognizes shortened poly(A) tails, and decapping enzyme Dcp2 (Charenton et al., 2017). Hence, coupling of deadenylation and decapping in 5’-to-3’ mRNA decay involves formation of protein bridges between mRNA ends. Binding of these protein bridges to mRNA ends is likely stabilized by the intrinsic closeness of mRNA ends.

### 3.3. mRNA compactness and the closed-loop model of translation initiation

Another example of protein-mediated interactions between the 5’ cap and 3’ poly(A) tail is the formation of closed-loop structure during translation initiation. To begin protein synthesis, a complex containing the small ribosomal subunit and a number of initiation factors is recruited to the m^7^Gppp cap structure at the 5’ end of the mRNA (Sonenberg, 2008). The cap structure is recognized by initiation factor <u>eIF4E</u> (<u>e</u>ukaryotic Initiation <u>F</u>actor 4E). A complex, called eIF4F, which is formed by eIF4E and two other initiation factors eIF4G and eIF4A, recruits the small ribosomal subunit preassembled with initiator tRNA and initiation factors 1, 1A, 2, 3 and 5 (Aitken & Lorsch, 2012). After recruitment to the 5’ end of mRNA, the small (40S) ribosomal subunit is believed to scan the 5’ UTR of mRNA until it reaches the start codon.

The poly(A) tail at the 3’ end of mRNA was shown to stimulate translation initiation (Jacobson & Favreau, 1983; Munroe & Jacobson, 1990). A number of studies showed that stimulation of translation by the combination of a cap and a poly(A) tail is greater than the product of stimulatory effects of a cap and a poly(A) tail alone (Thompson & Gilbert, 2017). This phenomenon was described as “synergy” between the 5’ cap and 3’ poly(A) tail (Gallie, 1991). The cap-poly(A) tail synergy is believed to be mediated by the binding of cap-binding factor eIF4E and poly(A) binding protein (PABP) to different parts of eIF4G (Kahvejian, Svitkin, Sukarieh, M’Boutchou, & Sonenberg, 2005; Tarun & Sachs, 1996; Tarun, Wells, Deardorff, & Sachs, 1997). The eIF4E•eIF4G•PABP complex was thought to “circularize” the mRNA (making a “closed loop”) (Jacobson, 1996; Wells, Hillner, Vale, & Sachs, 1998). Mounting evidence suggests that mRNAs vary in the degree to which their translation depends on the presence of eIF4G-PABP interactions (Amrani, Ghosh, Mangus, & Jacobson, 2008; Arava et al., 2003; Archer, Shirokikh, Beilharz, & Preiss, 2016; Thompson & Gilbert, 2017; Thompson, Rojas-Duran, Gangaramani, & Gilbert, 2016). Nevertheless, the interaction between eIF4E, eIF4G and PABP is conserved from yeast to humans and is thought to play an important role in the initiation of protein synthesis in eukaryotes.

Contrary to the idea of eIF4E•eIF4G•PABP-driven circularization of mRNA, the evidence suggests that in cells, intramolecular basepairing of the mRNA is bringing together the 3’ end of the 3’ UTR and the 5’ end of mRNA rather than the eIF4E•eIF4G•PABP complex. One line of evidence is that the eIF4E•eIF4G complex was found to crosslink to the 3’ end the 3’ UTR of yeast transcripts in cells without PABP (Archer, Shirokikh, Hallwirth, Beilharz, & Preiss, 2015). Another line of evidence comes from single-molecule-resolution fluorescent *in situ* hybridization (smFISH) imaging studies in human cells. The median distance between the 5’ and 3’ ends of actively translated mRNAs, with lengths from 6,000 to 18,000 nucleotides, was 100-200 nm (Adivarahan et al., 2018; Khong & Parker, 2018). By contrast, the 5’ and 3’ ends in these mRNAs co-localized when translation was inhibited with arsenite or the antibiotic puromycin (Adivarahan et al., 2018; Khong & Parker, 2018). Strikingly, the colocalization of mRNA ends was observed in cells in which the eIF4G•PABP interaction was disrupted by mutagenesis (Adivarahan et al., 2018). Based on these experiments, it was concluded that, in the absence of helicase activity of translating ribosomes, mRNA ends come in close proximity because of intramolecular basepairing interactions within mRNA (Adivarahan et al., 2018; Khong & Parker, 2018). Hence, at least in the translationally repressed state of mRNA, for example, in mRNAs sequestered into stress granules (Adivarahan et al., 2018; Khong & Parker, 2018), mRNA ends are brought in close proximity by mRNA secondary structure. These studies also indicated that the cap•eIF4E•eIF4G•PABP•poly(A) complex is not constitutively bound to actively translated mRNAs. Hence, the cap•eIF4E•eIF4G•PABP•poly(A) interactions may be transient and form during the transition from a translationally inactive to translationally active state of mRNA.

By bringing the 5’ and 3’ ends in the proximity of a few nm, mRNA intramolecular secondary structure decreases the entropic penalty for the cap•eIF4E•eIF4G•PABP•poly(A) complex formation. In other words, PABP binding to the poly(A) tail can facilitate the recruitment of the cap-binding protein complex, eIF4E•eIF4G, to the 5’ end of the mRNA since the 5’ and 3’ ends of mRNA are intrinsically close. Hence, the eIF4G-PABP interaction may have emerged throughout evolution to exploit the intrinsic closeness of mRNA ends.

### 3.4. The intrinsic closeness of mRNAs ends may facilitate regulation of translation initiation

**\**While the role of the intrinsic closeness of mRNA ends in the cap•eIF4E•eIF4G•PABP•poly(A) complex formation needs to be further examined, involvement of basepairing interactions between the 5’ and 3’ ends of mRNA in translation initiation of positive-strand RNA plant viruses has been demonstrated experimentally. Translation of these mRNAs, which lack the 5’ cap and poly(A) tail, requires the 3’ Cap-Independent Translational Enhancers (CITEs) sequence at the 3’ end of viral transcripts (Nicholson & White, 2011). The 3’ CITEs bind eIF4E•eIF4G (or eIF4G alone) and then recruit these initiation factors to the 5’ end of viral transcripts in the process that involves basepairing interactions between the 3’ CITE and the 5’ UTR of the same transcript (Nicholson & White, 2011).

A similar mechanism may underlie translation of histone mRNAs, which also lack the poly(A) tail. During translation initiation on histone mRNAs, the 40S subunit was shown to be initially recruited to specific structural RNA elements within the ORF and at the 3’ end of the mRNA (Cakmakci, Lerner, Wagner, Zheng, & Marzluff, 2008; Martin et al., 2011). The subsequent binding of the 40S to the start codon near the 5’ end of the transcript is likely facilitated by the intrinsic compactness of the mRNA.

The intrinsic compactness of mRNA may also underlie numerous examples of message-specific translational control of protein expression mediated by the formation of protein bridges between the 5’ cap and specific regulatory sequences in the 3’ UTR. For example, translation of mRNAs containing a cytoplasmic polyadenylation element (CPE) in the 3’ UTR is repressed by recruitment CPEB•Maskin protein complex to CPE. CPEB•Maskin binds to eIF4E•5’ cap and displaces eIF4G (Sonenberg & Hinnebusch, 2009). Similarly, translation of *Drosophila oskar* mRNA is inhibited by the binding of the eIF4E•5’ cap to the Bruno•Cup protein complex tethered to Bruno response element (BRE) in the 3’ UTR of *oskar* mRNA (Nakamura, Sato, & Hanyu-Nakamura, 2004). Translational repression of ceruloplasmin mRNA upon interferon-γ treatment is mediated by formation of the GAIT complex assembled from Glu-Pro-tRNA synthetase, NS-associated protein 1, GAPDH, and 60S ribosomal protein L13a, which is released from the 60S subunit by phosphorylation. The GAIT complex simultaneously binds to the GAIT element in the 3′ UTR and eIF4G at the 5’ end of ceruloplasmin mRNA to block recruitment of the small ribosomal subunit (Arif et al., 2018).

## Conclusions

Short end-to-end distances might facilitate the binding of protein factors that regulate translation initiation and mRNA decay by bridging mRNA 5’ and 3’ ends. There has been a long-standing view that protein-protein interactions bring the two ends of mRNAs in proximity. This review focuses on the recent understanding that RNA sequences fold by basepairing to bring the 5’ and 3’ ends in proximity, based on computation and on FRET measurements. Closeness of RNA ends is a property of most, if not all, mRNAs and lncRNAs. Only sequences that are devoid of guanosines and have low sequence complexity are intrinsically unstructured. Alternatively, RNA ends can be kept apart by pseudoknotted basepairing interactions with nucleotides in the middle of the RNA. For example, a pseudoknot in the small subunit rRNA facilitates the formation of a structure that makes the 5’ and 3’ ends distant and also exposes the anti-Shine-Dalgarno sequence for basepairing.

Given the intrinsic propensity of mRNA to fold into structures with short end-to-end distances, it makes sense that proteins would have evolved to complexes that specifically recognize both ends of mRNA. The prevalence of these protein bridges between the 5’ cap and 3’ UTR in the evolution of translational control supports the hypothesis that the regulatory protein complexes emerged throughout evolution to utilize the intrinsic compactness of mRNA.

